# SARS-CoV-2 omicron variants succumb *in vitro* to *Artemisia annua* hot water extracts

**DOI:** 10.1101/2022.07.22.501141

**Authors:** M.S. Nair, Y. Huang, P.J. Weathers

**Affiliations:** Aaron Diamond AIDS Research Center, Columbia University Irving Medical Center, New York, NY, USA; Department of Biology and Biotechnology, Worcester Polytechnic Institute, Worcester, MA 01609, USA

**Keywords:** *Artemisia annua*, tea infusions, Omicron, COVID-19, SARS-CoV-2, WA1, BA.1.1.529+R346K, BA.2, BA.2.12.1, BA.4

## Abstract

The SARS-CoV-2 (COVID-19) global pandemic continuous to infect and kill millions while rapidly evolving new variants that are more transmissible and evading vaccine-elicited antibodies. *Artemisia annua* L. extracts have shown potency against all previously tested variants. Here we further queried extract efficacy against omicron and its recent subvariants. Using Vero E6 cells, we measured the *in vitro* efficacy (IC_50_) of stored (frozen) dried-leaf hot-water *A. annua* L. extracts of four cultivars (A3, BUR, MED, and SAM) against SARS-CoV-2 variants: original WA1 (WT), BA.1.1.529+R346K (omicron), BA.2, BA.2.12.1, and BA.4. IC_50_ values normalized to the extract artemisinin (ART) content ranged from 0.5-16.5 µM ART. When normalized to dry mass of the extracted *A. annua* leaves, values ranged from 20-106 µg. Although IC_50_ values for these new variants are slightly higher than those reported for previously tested variants, they were within limits of assay variation. There was no measurable loss of cell viability at leaf dry weights ≤50 µg of any cultivar extract. Results continue to indicate that oral consumption of *A. annua* hot-water extracts (tea infusions) could potentially provide a cost-effective approach to help stave off this pandemic virus and its rapidly evolving variants.

## 1. Introduction

The novel coronavirus, SARS-CoV-2, with its rapidly evolving variants continues to plague the global population with nearly 600 million cases and >6 million deaths (https://coronavirus.jhu.edu/map.html, accessed June 26, 2022). The past year the omicron (B.1.1.529) variant of concern (VOC) emerged along with a number of subvariants, especially BA.4 and BA.5 [1]. These are highly transmissible (Omicron RO ≥ 10, Delta RO = 7 [2]) infecting even vaccinated individuals, albeit with less severe outcomes (https://www.healthdata.org/covid/COVID-19-vaccine-efficacy-summary, accessed June 26, 2022).

Omicron (B.1.1.529) and its B.1.1.529+R346K isolate have shown resistance to neutralization by antibodies in patients who have had COVID-19 or been vaccinated and even boosted with one of the widely used vaccines [3–5]. Recently, variants, BA.2.12.1 and BA.4/5 were shown to be 1.8 and 4.2 times, respectively, more resistant to sera from individuals who were vaccinated and boosted [6]. Additionally, recent clinical case studies showed that vaccinated and boosted individuals who took a course of Paxlovid™ have shown relapse and post relapse can accidently infect family members [7]. This presents an even more pressing need for an expanded diversification of therapeutics, which may also serve as prophylactics in a population setting.

Although a number of different drugs have been trialed [8] there are few approved small molecule drugs available to treat COVID-19. The antiviral drug Paxlovid™ was recently approved as a *per os* combination drug of nirmatrelvir (or PF-07321332) with ritonavir, developed by Pfizer with good anti-SARS-CoV-2 efficacy and relatively few adverse drug reactions [9,10]. Nirmatrelvir acts as a main viral protease inhibitor [9] with ritonavir enhancing its pharmacokinetics by inhibiting hepatic CYP3A4 [11]. Despite this success, access may be limited (https://www.nature.com/articles/d41586-022-00919-5, accessed June 26, 2022). While generic production is estimated at ≤25 USD, it also may be unaffordable to many, especially in low and middle income countries (https://www.reuters.com/business/healthcare-pharmaceuticals/generic-drugmakers-sell-pfizers-paxlovid-25-or-less-low-income-countries-2022-05-12/, accessed June 26, 2022). Thus, there remains a need for more cost-effective therapeutics to treat the global population.

Previously we and others showed that extracts of dried leaves of many cultivars of the medicinal plants, *Artemisia annua* L., which produces the antimalarial sesquiterpene lactone artemisinin (ART; Figure 1), and *A. afra* Jacq. ex Willd., a related perennial species lacking artemisinin, prevented SARS-CoV-2 replication *in vitro* [12–15]. Although ART has some anti-SARS-CoV-2 activity, we showed that antiviral efficacy was inversely correlated to ART content [12]. Here we report *in vitro* efficacy for four of the seven originally studied *A. annua* L. cultivars against omicron (B.1.1.529) and three of its subvariants: BA.2, BA.2.12.1, and BA.4.

**Figure 1.**
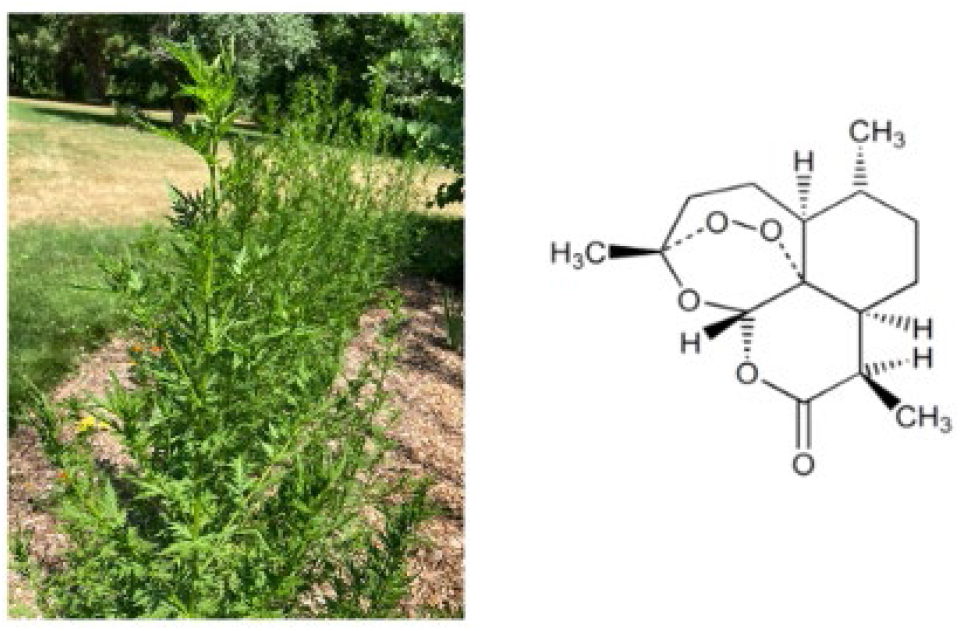
*Artemisia annua* L. and artemisinin.

## 2. Results and Discussion

Hot-water extracts of four cultivars of *A. annua* inhibited the recently evolved omicron and its three tested subvariants of SARS-CoV-2 with IC_50_ values calculated and normalized to the ART content of each tested tea infusion ranging from 0.5-16.5 µM ART (Figure 2; the lower the IC_50_, the more potent the drug/extract). When the IC_50_ values were instead normalized to the dry mass of the extracted *A. annua* leaves, values ranged from 20-106 µg (Figure 3). Although the values for these new variants are for the most part slightly higher than the IC_50_ values reported for variants previously reported [12,13] and all are summarized in Table 1, they fell within limits of assay variation. As already reported for extracts used in this study, there was no measurable loss of cell viability at a dry weight of ≤50 μg for any cultivar extract [12]. Others have reported *in vitro* efficacy of *A. annua* [14,15] and *A. afra* [14] extracts against earlier variants of SARS-CoV-2; however, to our knowledge there are no reports showing efficacy against omicron or its variants.

**Table 1.** Comparative IC50 values of *A. annua* L. hot-water extracts (10 g/L) against all tested strains of SARS-CoV-2 based on either artemisinin content or leaf dry weight (DW).

| Cultivar | Potency normalized to artemisinin content |  |  |  |  |  |  |  |  |  |  |
| --- | --- | --- | --- | --- | --- | --- | --- | --- | --- | --- | --- |
| | IC<sub>50</sub> μM artemisinin | | | | | | | | | | |
|  | <u>WA1*</u><br>(WT) | <u>B.1.1.7*</u><br>(alpha) | <u>B.1.351*</u><br>(beta) | <u>P.1**</u><br>(gamma) | <u>B.1.617.1**</u><br>(kappa) | <u>B.1.617.2**</u><br>(delta) | <u>WA1</u><br>(WT) | <u>BA.1.1.529+R346K</u><br>(omicron) | <u>BA.2</u> | <u>BA.2.12.1</u> | <u>BA.4</u> |
| SAM | 3.4 | 4.9 | 8.4 | 7.9 | 7.0 | 7.0 | 4.9 | 8.2 | 7.6 | 5.3 | 16.5 |
| A3 | 0.8 | 1.1 | 2.0 | 1.9 | 1.9 | 2.1 | 1.2 | 1.8 | 1.0 | 1.5 | 2.6 |
| BUR | 0.4 | 0.3 | 0.8 | 1.2 | 1.1 | 1.2 | 0.7 | 1.2 | 0.6 | 0.5 | 1.1 |
| MED | 2.9 | 2.0 | 3.6 | 2.9 | 2.5 | 4.8 | 4.2 | 4.7 | 2.1 | 3.9 | 6.4 |
| Cultivar | Potency normalized to dry mass of leaves used in tea infusion |  |  |  |  |  |  |  |  |  |  |
| | IC<sub>50</sub> μg leaf DW | | | | | | | | | | |
|  | <u>WA1*</u> | <u>B.1.1.7*</u> | <u>B.1.351*</u> | <u>P.1**</u> | <u>B.1.617.1**</u> | <u>B.1.617.2**</u> | <u>WA1</u> | <u>BA.1.1.529+R346K</u> | <u>BA.2</u> | <u>BA.2.12.1</u> | <u>BA.4</u> |
| SAM | 21.5 | 31.3 | 53.7 | 50.7 | 45.0 | 45.1 | 31.4 | 52.5 | 48.9 | 34.1 | 106.0 |
| A3 | 15.7 | 22.1 | 39.6 | 38.2 | 37.0 | 42.4 | 23.9 | 36.7 | 20.0 | 30.7 | 50.9 |
| BUR | 15.1 | 11.0 | 32.5 | 50.1 | 44.7 | 49.8 | 27.5 | 48.9 | 23.6 | 22.7 | 45.2 |
| MED | 41.7 | 28.2 | 51.5 | 41.0 | 37.0 | 67.7 | 59.7 | 67.0 | 30.0 | 56.0 | 90.6 |

\* Values taken from Nair et al. [12]; \*\* values taken from Nair et al. [13].
IC<sub>50</sub>s are values where virus is 50% inhibited. Data are an average of three replicates.

**Figure 2.**
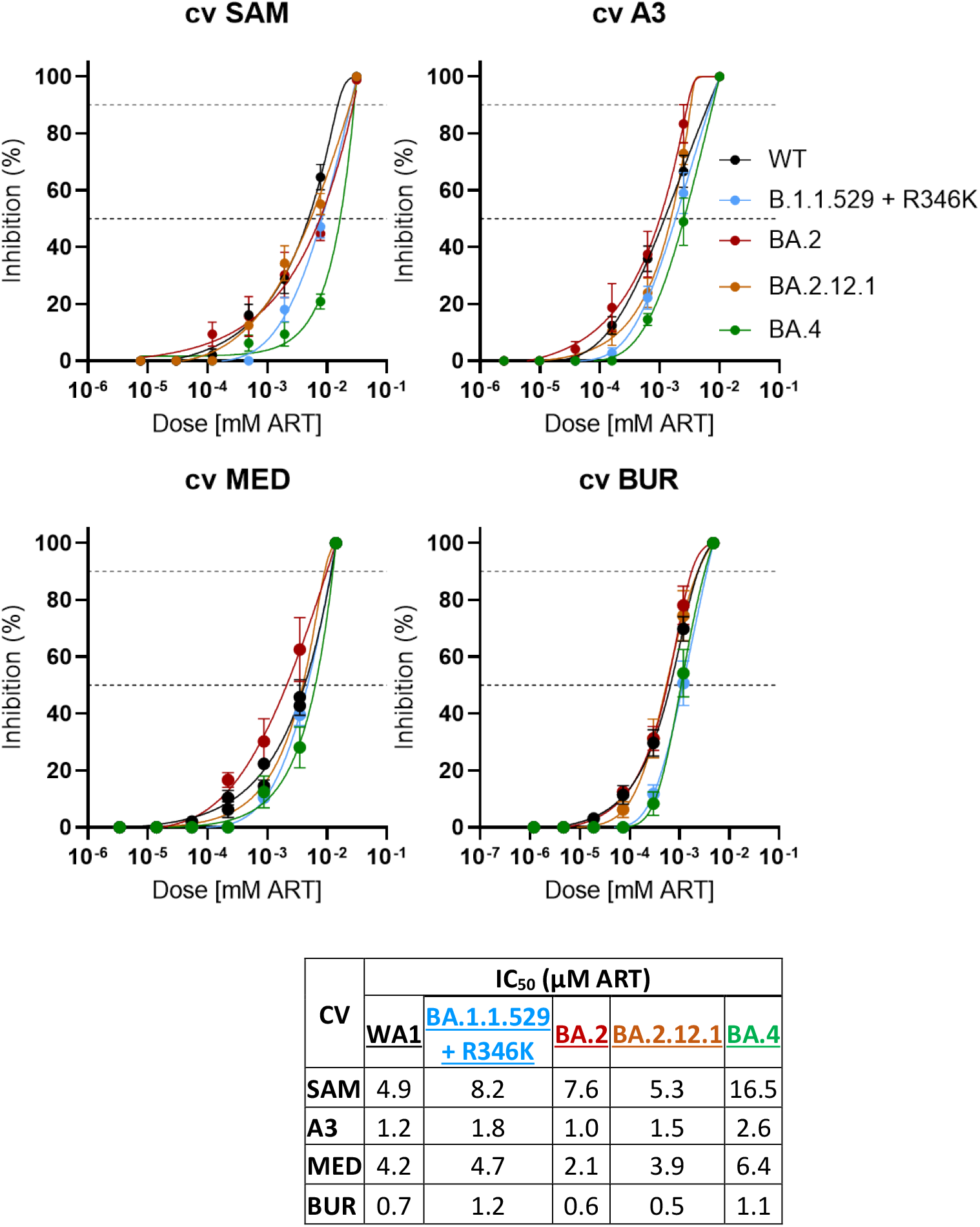
SARS-CoV-2 variant inhibition by four cultivars of *A. annua* L. hot water extracts normalized to their artemisinin content and compared to WT. WT, USA/WA1; variants: BA.1.1.529+R346K, omicron; omicron subvariants: BA.2; BA.2.12.1, and BA.4 at a multiplicity of infection (MOI) of 0.1 in Vero E6 cells. Data are plotted from an average of three replicates from each of two experiments ± SE.

**Figure 3.**
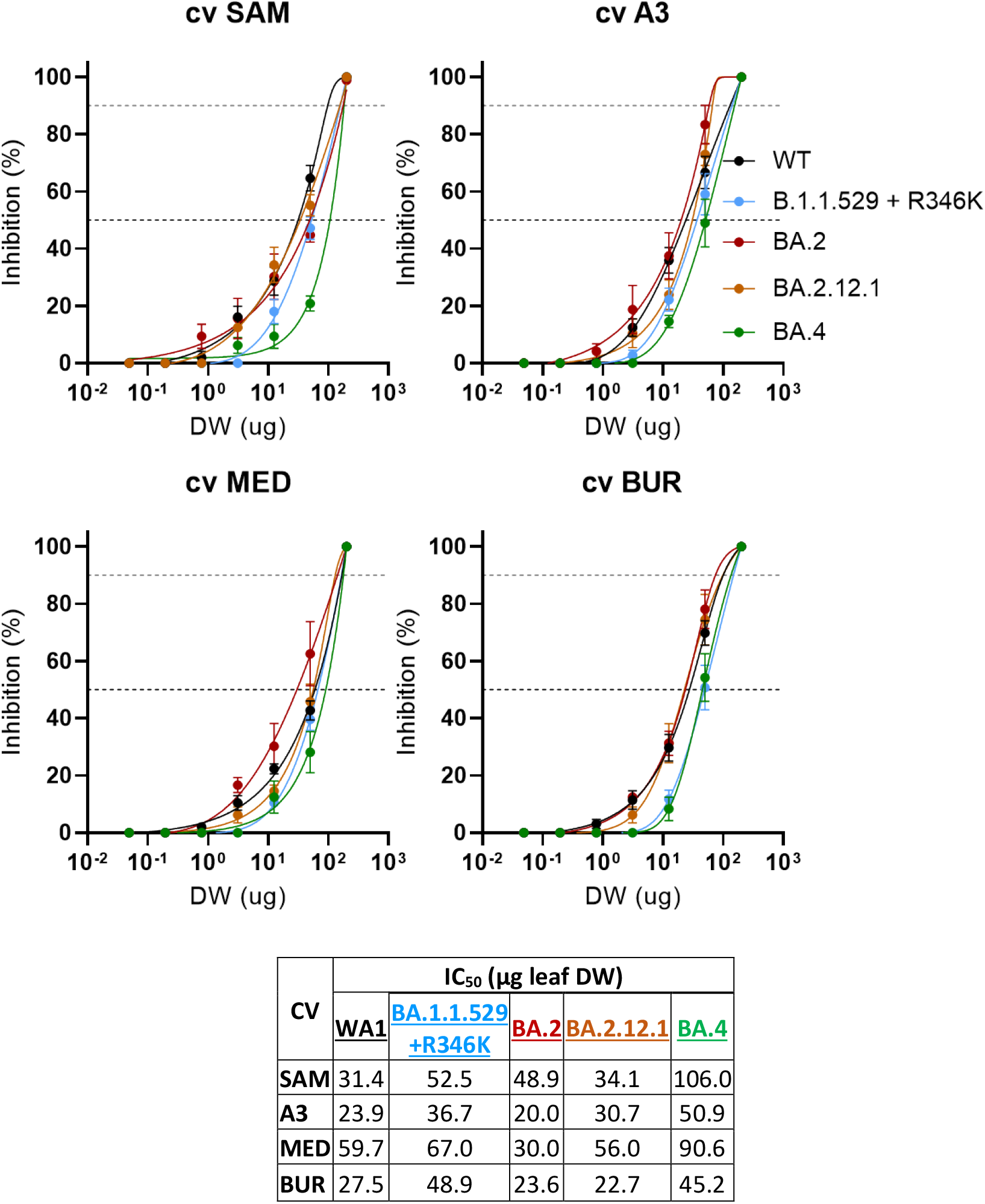
SARS-CoV-2 variant inhibition by four cultivars of *A. annua* L. hot water extracts normalized to their *A. annua* leaf dry mass (DW) and compared to WT. WT, USA/WA1; variants: BA.1.1.529+R346K, omicron; omicron subvariants: BA.2; BA.2.12.1, and BA.4 at a multiplicity of infection (MOI) of 0.1 in Vero E6 cells. Data are plotted from an average of three replicates from each of two experiments ± SE.

Although ART IC_50_ values are shown in this study, we previously reported that potency was inversely related to ART concentration [12] and others showed that *A. afra*, a species lacking ART, was also highly effective *in vitro* against SARS-CoV-2 [14]. Nevertheless, ART has some anti-SARS-CoV-2 activity as we and others showed [12,14–19]. Although there have been some clinical studies using ART, it was used as a combination therapy. For example, in a small non-randomized controlled trial where patients were treated with ART-piperaquine (ART-PPQ) or placebo the mean time for recovery where there was no longer PCR-detectable virus was 10.6 d for ART-PPQ treated patients vs. 19.3 d for those receiving placebo [20]. All patients treated with ART-PPQ were virus-free after 21 d compared to 36 d for placebo. In another small trial patients had faster recovery vs. placebo in those who used ArtemiC, an oral spray containing ART, curcumin, frankincense, and vitamin C [21]. To our knowledge, however, there are no reports of clinical trials using *A. annua* or its extracts.

Because we and others [12,14] showed that ART is not the most likely anti-SARS-CoV-2 therapeutic phytochemical in *A. annua* extracts, questions remain regarding the identity of these non-ART phytochemicals. To resolve that question, several groups have used *in silico* approaches [22,23]. Tang et al. screened the Traditional Chinese Medicines for systems Pharmacology Databased and Analysis Platform to identify all phytochemicals reportedly in *A. annua* then ranked them according to oral bioavailability and drug likeness (OB and DL, respectively). That list was narrowed to 19 compounds within their OB and DL limits of ≥30% and 0.18, respectively. They concluded that many on the list of 19 compounds had anti-inflammatory, immune regulatory, and therapeutic properties. Among the top therapeutic candidates were luteolin and isorhamnetin. Using a ZINC library the Efferth lab also screened an *in silico* library of >39,000 natural product compounds including some from plants with known antiviral activity and narrowed their hits to 33 likely compounds [22]. Of the top 12, three, isorhamnetin, luteolin, and rosmarinic acid, are present in *A. annua* and when tested *in vitro* had IC_50_ values of 8.42, 11.81, and 9.43 µM, respectively, against the main protease in SARS-CoV-2, 3CL^pro^, a chymotrypsin-like protease involved in viral replication. Along with reports of anti-SARS-CoV-2 activity of other *A. annua* phytochemicals, e.g., quercetin and myricetin against NTPase/helicase [24,25], many other small molecules, especially flavonoids, are showing antiviral potential and likely work in combination (synergistically) in these extracts to achieve the therapeutic response. Future studies are needed to identify and validate the activity of these anti-SARS-CoV-2 therapeutic compounds.

## 3. Materials and methods

### 3.1 Plant material, extract preparations, and artemisinin analyses

Hot-water extracts (tea infusions) were previously prepared from dried leaves of *Artemisia annua* L. (SAM, MASS 00317314; BUR, LG0019527; A3, Anamed; MED, KL/015/6407) In brief: 10 g dried leaves/L were boiled in water for 10 min, solids removed via sieving, then 0.22 µm filter-sterilized and stored at - 80°C for this study and as detailed in Nair et al. [12]. ART analyses of tea infusions were by gas chromatography-mass spectrometry and detailed in Martini et al. [26] with ART contents detailed in [12]: ART in µg/mL were: 42.5 for A3; 20.1 for BUR; 59.4 for MED; and 149.4 for SAM.

### 3.2 Viral culture and infection

Cultivation of Vero E6 cells (ATCC CRL-1586) and viral infection are detailed in [12]. SARS-CoV-2 isolates (Table 2) were sourced from BEI Resources (www.beiresources.org). To determine their tissue culture infectious dose (TCID), viruses were titrated after propagation in Vero E6 cells, aliquoted, and frozen at - 80°C until later use. Multiplicity of infection (MOI) was 0.1 as used in other studies [27].

**Table 2.** SARS-CoV-2 isolates used in this study.

| SARS-CoV-2 isolate | BEI Resource Catalogue number |
| --- | --- |
| USA/WA12020 | NR-52281 SARS-Related Coronavirus 2, Isolate USA-WA1/2020 |
| Omicron B.1.1.529 + 346K | NR-56475 SARS-Related Coronavirus 2, Isolate hCoV-19/USA/HI-CDC-4359259-001/2021 |
| Omicron BA.2 | NR-56520 SARS-Related Coronavirus 2, Isolate hCoV-19/USA/CO-CDPHE-2102544747/2021 |
| Omicron BA.2.12.1 | NR-56781 SARS-Related Coronavirus 2, Isolate hCoV-19/USA/NY-MSHSPSP-PV56475/2022 |
| Omicron BA.4 | NR-56806 SARS-Related Coronavirus-2, Isolate hCoV-19/USA/MD-HP30386/2022 |

### 3.3 Drug inhibition assays of SARS-CoV-2 and cell viability

Dilutions of extracts were incubated for 1 h in 96-well tissue culture plates having a monolayer of Vero E6 cells seeded the prior day at 20,000 cells/well. SARS-CoV-2 virus was added to each well one h after extract addition to a final MOI of 0.1. Cells were cultured for 3 days in 5% CO_2_ at 37°C and then scored for cytopathic effects as previously detailed [27] and values converted into percent of control. Drug concentrations were log transformed and the concentration of drug(s) that inhibited virus by 50% (*i.e*., IC_50_), and the concentration of drug(s) that killed 50% of cells (*i.e*., CC_50_; viability), were log transformed and determined via nonlinear logistic regressions of log(inhibitor) versus response-variable dose-response functions (four parameters) constrained to a zero-bottom asymptote by statistical analysis. We already reported viability of Vero E6 cells post extract treatment in Nair et al. [12] for the same extracts. To normalize the IC_50_ values for the new variants tested or the WT and variants tested previously, dry mass of leaves and total ART contents measured in the *Artemisia* extracts were used as reported in Nair et al. [12].

### 3.4 Chemicals and reagents

Reagents were procured from Sigma-Aldrich (St. Louis, MO). Renilla-Glo was from Promega (E2720). EMEM (Cat # 30-2003) and XTT reagent (Cat # 30-1011k) were from ATCC.

### 3.5 Statistical analyses

The anti-SARS-CoV-2 analyses were done at least in triplicate. Plant hot water extracts had n≥6 independent assays as documented in Nair et al. [12]. IC_50_ values were calculated using GraphPad Prism V9.3.

## 4. Conclusions

Hot-water (tea infusion) extracts of *A. annua* continue to show activity against SARS-CoV-2 and the newest VOCs including omicron and three of its highly transmissible subvariants. Although the specific phytochemicals have not yet been identified, there are a number of possible candidates emerging in the literature. Validation of *A. annua* extracts against SARS-CoV-2 VOCs in a rodent model are needed as a next step towards human trials. Nevertheless, this plant is safe to use, and we urge testing in clinical trials sooner rather than later. WHO announced in 2021 that through its COVID-19 Solidarity Therapeutics Plus Trial that it has included intravenous artesunate as one of three repurposed drugs to treat COVID-19 [28]. Results are not anticipated until 2023, (last accessed July 11, 2022, https://www.isrctn.com/ISRCTN18066414). Regardless of outcome, and based on the continuing efficacy of *A. annua* extracts against all tested variants (10 to date), we again urge the WHO to consider including encapsulated dried leaf *A. annua* as a separate arm in their trial. Use of *A. annua* does not induce ART resistance [29], but the plant could be crucial in helping many in the world where access to vaccines and standard therapeutics is logistically challenging.

## Author Contributions

Conceptualization, M.S.N., P.J.W.; methodology, M.S.N., Y.H.; formal analysis, M.S.N., Y.H.; writing—original draft preparation, P.J.W.; writing—review and editing, M.S.N. and P.J.W.; supervision, M.S.N. All authors have read and agreed to the published version of the manuscript.

## Institutional Review Board Statement

Not applicable.

## Informed Consent Statement

Not applicable.

## Data Availability Statement

Data are contained within the article.

## Acknowledgments

We thank Prof. David Ho of the Aaron Diamond AIDS Research Center at Columbia University for supporting the live virus work in his lab. Prof. David Fidock, Columbia University, and Dr. Melissa Towler, Worcester Polytechnic Institute, provided critical review of the manuscript.

